# Retinotopic mapping data permit accurate matching of participants across different datasets

**DOI:** 10.1101/2024.12.19.629552

**Authors:** D. Samuel Schwarzkopf, Catherine A. Morgan, Reece P. Roberts, Steven C. Dakin

**Affiliations:** School of Optometry & Vision Science, University of Auckland, New Zealand; Experimental Psychology, University College London, United Kingdom; School of Psychology and Centre for Brain Research, University of Auckland, New Zealand; Centre for Advanced MRI, UniServices Limited, Auckland, New Zealand; Institute of Ophthalmology, University College London, United Kingdom

## Abstract

Public sharing of neuroimaging data is becoming increasingly common for the advancement and validation of scientific research. However, this sharing poses challenges regarding privacy and data safety, and associated questions about the ownership of research data. Here we show that a simple pattern correlation algorithm can match nominally deidentified participants across two separate functional magnetic resonance imaging (fMRI) experiments. This re-identification procedure is effective despite functional maps being spatially warped to a common template brain. This work highlights the need for appropriate safeguards against possible misuse of shared neuroimaging data.

## Introduction

Large and openly accessible neuroimaging datasets like the human connectome project and the THINGS dataset^1–3^ are a useful tool for advancing our understanding of the brain and for ensuring reproducibility of cognitive neuroscience research. However, it is crucial to protect participants’ identity. Brain data acquired with techniques like functional magnetic resonance imaging (fMRI) are unique. Informed consent processes usually explain to participants that their data are “deidentified” (pseudonymized) by assigning unique identifiers only known to the researchers immediately involved in the study, and that there is no way for others to trace their data back to the individual participant. But how true is this when it comes to functional brain maps? Do processing steps such as “defacing” structural brain images or only sharing normalized brain maps really make it impossible to reidentify a participant in a dataset?

Here, we tested the possibility of participant matching with typically used fMRI data. Previously, we conducted a small study to compare retinotopic mapping data of three participants scanned at two sites: a 3 Tesla scanner in Auckland, New Zealand, and a 1.5 Tesla scanner in London, United Kingdom^4^. The Auckland data were part of a larger study comprising 25 participants^5^. Our present analysis sought to match the maps of each London participant to their corresponding maps in the Auckland dataset, after they were spatially normalized to a common template brain (Figure 1).

**Figure 1.**
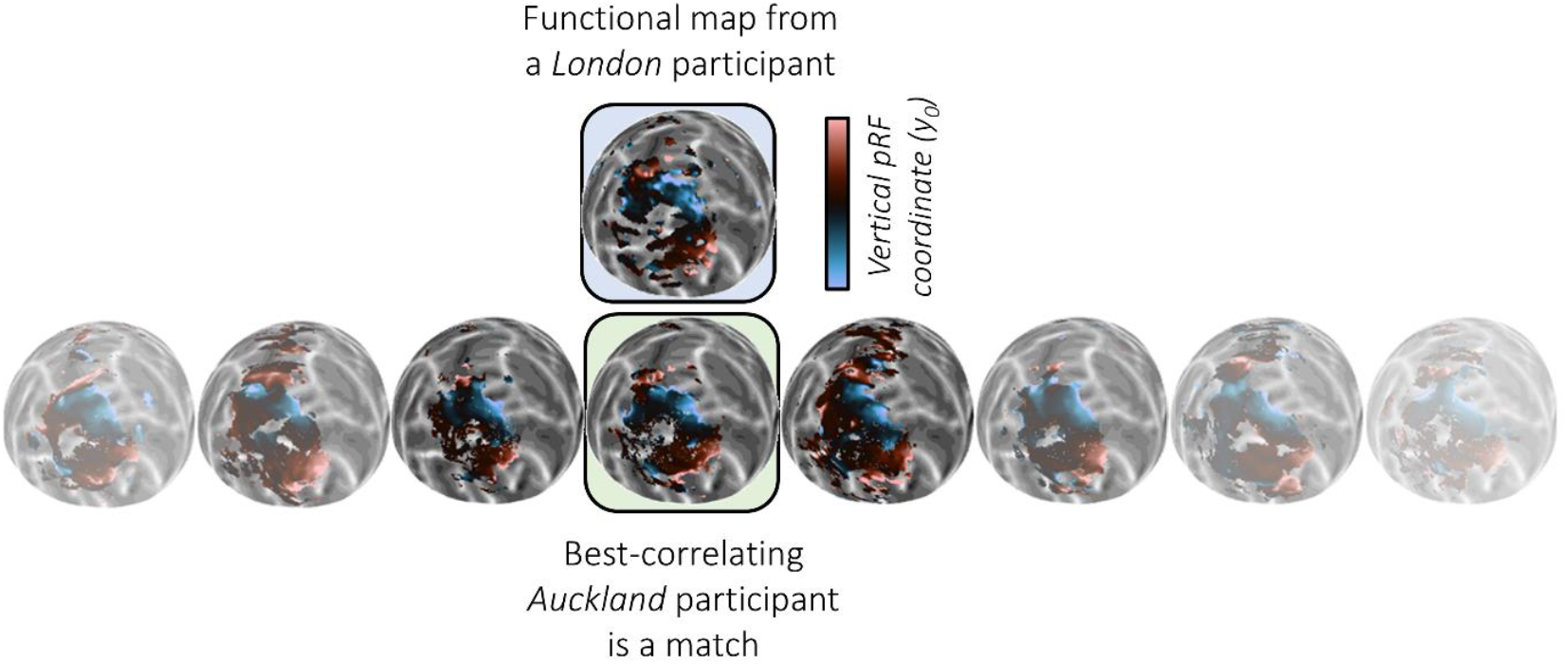
Procedure for matching participants between two data sets, based on their fMRI brain maps alone. Brain maps (here, y0 pRF coordinate) have been spatially normalized to a common template brain and shown on an inflated, spherical model of the occipital lobe. The map of a given London participant is compared to all the maps of Auckland participants. In this case, the Auckland participant whose map correlated best is indeed the one matching the London participant.

## Methods

### Participants

Twenty-five participants (15 females, 10 males; ages: 19– 47 years) were scanned at the Centre for Advanced Magnetic Resonance Imaging (CAMRI) at the University of Auckland, New Zealand. Their data were used to develop a new connective field mapping algorithm^5^. Three of these participants (2 females, 1 male; ages: 30-47 years) also participated in a retinotopic mapping experiment at the Birkbeck-University College London Centre for NeuroImaging (BUCNI) in London, United Kingdom. The purpose of these scans was to ascertain the similarity of population receptive field (pRF) measures between scanning sites^4^. The time between scans was several months for each participant. All procedures were approved by local ethics boards at each site.

### Stimuli

Inside the MRI scanner, participants viewed a screen (resolution: 1920 × 1080 pixels) positioned behind the scanner bore, so that it subtended 34° by 19° visual angle via a mirror mounted on the head coil. In Auckland, the screen was an MRI-compatible 32-inch liquid crystal display (BOLD Screen, Cambridge Research Systems Ltd.). In London, a custom-made screen was mounted inside the bore and stimuli were projected onto it.

Participants fixated a central blue dot target (diameter: 0.09°) while they viewed bar stimuli moving across the screen. Bars contained a black and white ripple pattern^6^. They traversed a circular region of the visual field, centered on fixation and extending out to an eccentricity of 9.5°. Stimulus contrast was ramped down at the outer edge of the circular region and immediately around the fixation target. Bars were 0.53° wide and moved in steps of 0.38°, one step every 1 s. Bars swept the visual field in eight directions, with each sweep lasting 25 s. After the fourth and eighth sweep, we presented a 25 s blank period without bar stimuli to estimate the baseline response. The order of bar sweep directions was identical in each scanning run.

### Procedure

Each participant was scanned with six runs of the pRF mapping experiment, with each run lasting 4 min 10 s. Participants were performing an incidental task monitoring the fixation target for a color change to purple. This event could occur every 200 ms with a probability of 0.01, with the constraint that events could not occur consecutively. At each site, we also collected a structural brain image in addition to the functional retinotopic mapping data.

### Scanning parameters

At the Auckland site, scans were conducted using a Siemens MAGNETOM Skyra 3 Tesla scanner. In London we used a Siemens MAGNETOM Avanto 1.5 Tesla scanner. At both sites, we used a 32-channel head coil with the frontal portion removed, which left 20 effective receive channels. We acquired blood oxygen level dependent signal using T2*-weighted images. At both sites, we used an accelerated multiband sequence^7,8^ at 2.3 mm isotropic voxel size, matrix size 96×96, 36 transverse slices angled to be approximately parallel to the calcarine sulcus, in a TR of 1000 ms. Site parameter variations were as follows: In Auckland, TE 30 ms, flip angle 62°, multiband/slice acceleration factor 3, in-plane/parallel imaging acceleration factor 2, rBW: 1680Hz/Px; In London, TE: 55 ms, flip angle: 75°, multiband/slice acceleration factor, 4, rBW: 1628 Hz/Px.

Moreover, we collected T1-weighted structural brain images at each site with the full 32-channel head coil. This was a magnetization-prepared rapid acquisition with gradient echo scan with 1 mm isotropic voxel size, matrix size 256 × 256, and full brain coverage. Site parameter variations in Auckland: flip angle: 8°, TI: 880 ms, TR: 2000 ms, TE: 2.8 ms, 208 sagittal slices; and in London: flip angle: 7°, TI: 1000 ms, TR: 2730 ms, TE: 3.57 ms, 176 sagittal slices.

### Data preprocessing

We reconstructed three-dimensional models of the cortical surface (pial and white-grey matter boundaries) from the structural brain image using FreeSurfer^9–11^. For the three participants scanned at both sites, we did this separately for the London and Auckland structural images. Next, functional images were preprocessed in SPM12 (https://www.fil.ion.ucl.ac.uk/spm/software/spm12) by applying motion correction and coregistration of the functional images to the structural scan, using default parameters. Finally, functional images were projected to the cortical surface by determining for each vertex in the cortical model the functional imaging voxel that fell halfway between the white-matter and pial boundaries.

Functional data from each participant and scan were then analyzed using population receptive field (pRF) mapping^5,12,13^, using our analysis toolbox SamSrf (https://osf.io/2rgsm). In short, this entails a forward model predicting the neural response of a two-dimensional Gaussian pRF with given parameters to the stimulus. We then convolved this predicted neural time series with a canonical hemodynamic response function based on a large normative dataset acquired previously^14^. The analysis determines the optimal pRF parameters to explain the observed fMRI time series for each vertex using a least squared fit. To save computation time, analyses were restricted to only the occipital cortex.

Our functional data contained four maps of primary variables for each participant^4^: the horizontal (x0) and vertical (y0) position of pRF centers, the pRF size (σ), and the goodness-of-fit (R^2^) of the pRF model. The pRF positions were further converted into polar coordinates, that is, polar angle and eccentricity maps. Next, we used a nearest neighbor algorithm to spatially normalize these parameter maps from native brain space to a common template brain (the standard *fsaverage* in FreeSurfer). Therefore, we now had maps in the same cortical space, 25 from Auckland and 3 from London.

## Results

We tested whether we could match the three London participants to their data in the larger Auckland dataset from their fMRI data alone. We calculated the Pearson correlation between the spatially normalized maps (vertex-wise patterns of values) for each London participant with all the 25 Auckland participants. We took the Auckland participant showing the strongest correlation for each map as the match for each of the three London participants (Figure 1). This simple procedure allowed perfect matching of each of the three participants for the R^2^, y0, and σ maps (Figure 2). Assuming a binomial probability of 1/25, this result is extremely unlikely by chance (p<0.0001, Bayes factor in favor of alternative hypothesis, BF10>3906). Interestingly, the identification was poorer for x0 maps and only identified one of three participants correctly, which was not significant (p=0.1153, BF10=2.26). Presumably because of this reduced accuracy for the x0 data, reidentification based on polar angle was also poorer (Figure 2E), although still significant (polar: p=0.0047, BF10>54). In contrast, eccentricity (Figure 2F) did not permit above chance reidentification (p=0.1153, BF10=2.26). Thus, it was presumably the coarse-scale maps related to visual responsiveness, upper-vs-lower visual field representation, and spatial selectivity that afforded the best capacity for reidentification, rather than the fine-grained retinotopic organization.

**Figure 2.**
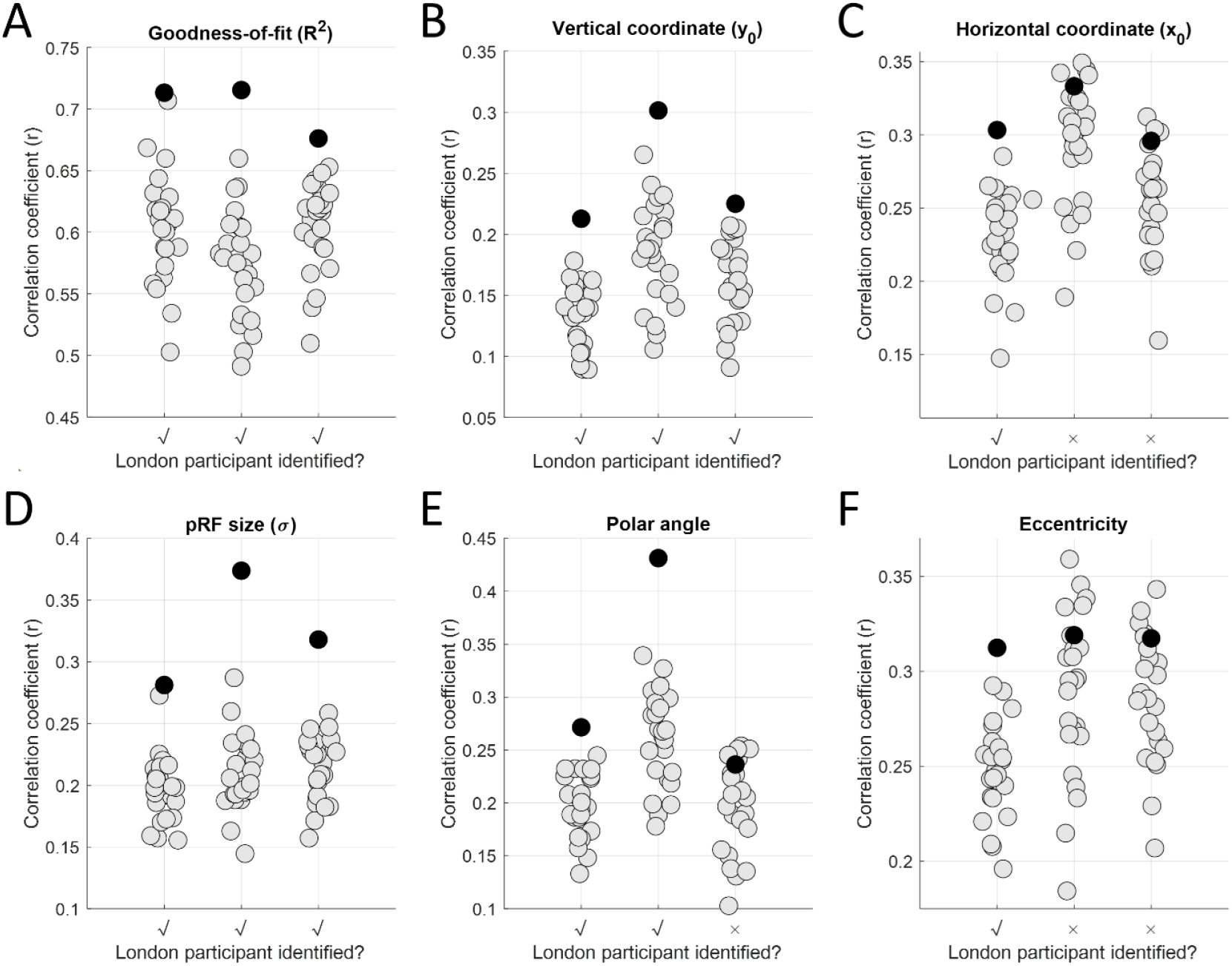
Reidentification of participants across datasets based on different fMRI parameters, specifically the goodness-of-fit (**A**), vertical coordinate (**B**), horizontal coordinate (**C**), pRF size (**D**), polar angle (**E**), and eccentricity (**F**). Symbols denote the correlation between each Auckland participant and each of the three London participants. Grey discs: non-matching participants. Black discs: matching participant. The x-axis indicates whether the London participant was correctly identified as the one with maximal correlation. Note that for polar angle maps we used circular correlation; all other maps were compared using Pearson correlation.

In this first analysis we used all vertices in the three-dimensional model mesh of the occipital cortex. That might have predisposed the analysis to detecting the broad activation pattern. In a secondary analysis, we restricted the pattern correlation to only those vertices in the brain model that consistently showed any visual response. This also yielded strong accuracy, although the pattern of results changed somewhat (Figure 3). While reidentification performance was less than perfect (2 of 3) for R^2^ and y0, it was still perfect for σ maps. Also, in contrast to the first analysis, accuracy was now perfect for x0 and eccentricity. Interestingly, polar angle maps did not permit accurate reidentification of any of the three London participants in this analysis.

**Figure 3.**
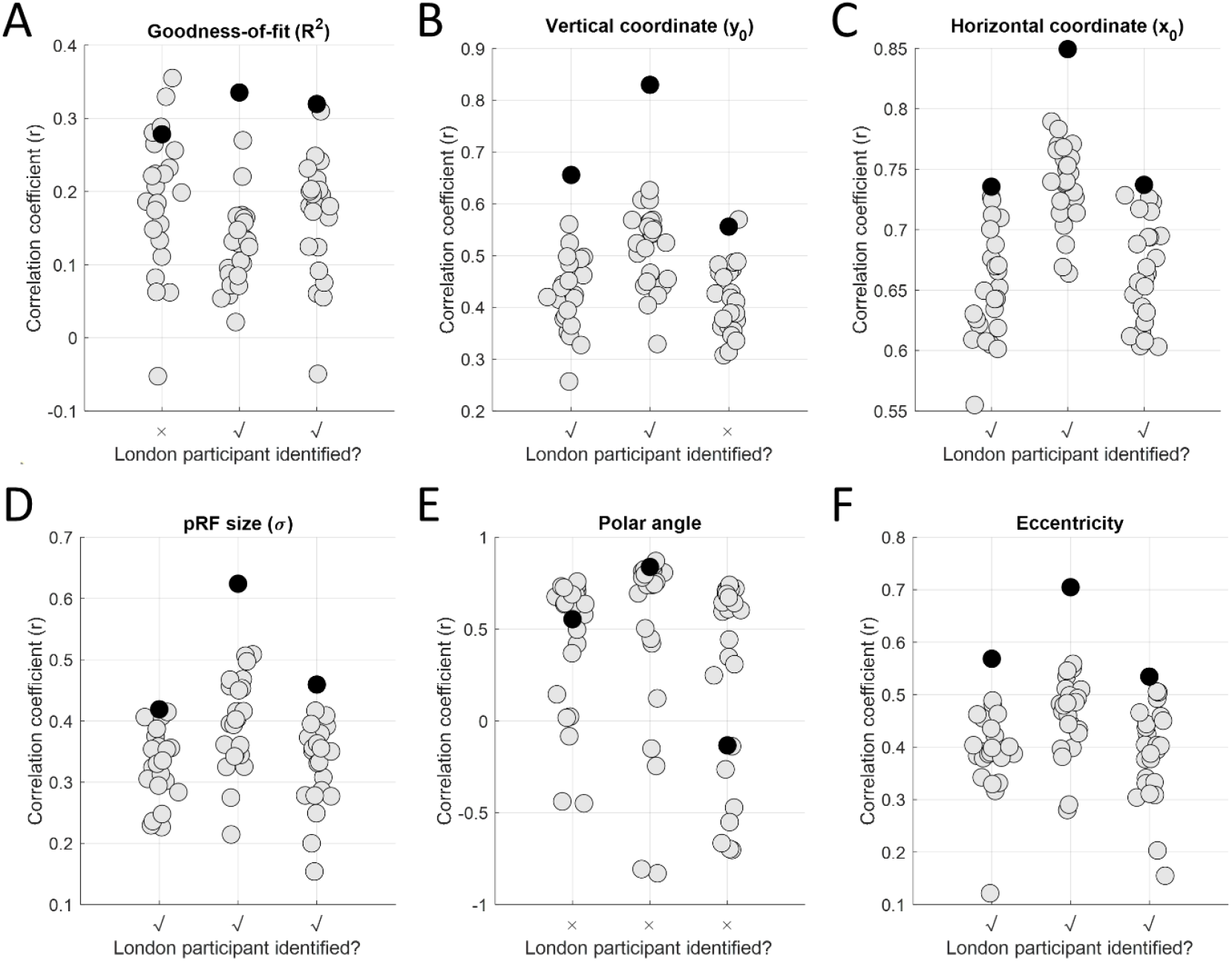
Reidentification of participants across datasets based on different fMRI parameters. All conventions as in Figure 2, except that here the correlations were calculated only for vertices where there was any visual response in both maps in a given pair.

While not all map types afforded perfect reidentification for all three London participants, combining the information across the different variables may help disambiguate participants. To test this, we calculated the average correlation across all six maps for each participant pair and repeated the analysis. This procedure correctly identified all three London participants for the whole occipital cortex data, but only two of the three when restricting the analysis to visually responsive cortex.

## Discussion

We show that it is straight-forward to match nominally deidentified fMRI data of participants across different datasets. For some map types, reidentification was perfect: correctly picking out each of the three participants scanned in London from the larger Auckland group. When considering the whole occipital cortex, this analysis was most accurate for coarse-scale patterns, presumably related to general visual responsiveness and spatial selectivity (pRF size). However, restricting this analysis to only parts of cortex that showed visual responses in each participant also yielded highly accurate reidentification for some of the finer-grained functional architecture, specifically the Cartesian coordinates, eccentricity, and pRF size parameters.

Critically, reidentification was possible even though functional maps had been spatially normalized to a common template brain. This may seem counterintuitive: normalization is designed to make individual data more alike. Should this therefore not reduce the capacity for identification? Spatial normalization brings different brains into alignment, based on common anatomical landmarks. The deviation at each point in the template space between the group mean and each individual brain is that individual’s unique signature. It is therefore possible that spatial normalization in fact boosts reidentification accuracy for functional data because it corrects for variations between scans.

Our algorithm used a simple pattern correlation to compare maps. This procedure relies on having a common brain space. However, more sophisticated methods based on machine learning might result in even higher accuracy to reidentify participants. It is conceivable that they can work using native brain space data. Indeed, native brain space might contain additional variables that could be informative for identification, such as brain size, variations in cortical thickness, or idiosyncratic gyrification. Our aim here was not to test the limits of what is possible. Rather, our study proves the concept that using standard analysis procedures for this type of data, without using any explicit anatomical information, can provide startlingly accurate matches.

While our sample is not particularly large, reidentifying all three London participants among the 25 Auckland maps is no mean feat. Such accuracy is extremely unlikely to occur by chance. However, reidentification accuracy was not perfect for all maps, and this depended on some analytical choices. Someone attempting such an analysis would therefore not necessarily know which parameters afford the best accuracy. But unfortunately, this does not diminish the risk. The real danger is not that every person in a large database of fMRI scans can be reidentified with high accuracy every time. An imperfect indication of a match could suffice to identify a person. Someone trying to match datasets could be an unethical actor targeting a particular individual and they may have access to additional information (such as demographic details) that further narrows down the options. More generally, combining multiple reidentification values and other available information could improve the match further. We showed basic proof of this concept using the average correlation values across the six map types. Here, again, a more sophisticated clustering algorithm and incorporating adjacent information probably yields even better reidentification accuracy.

This reidentification algorithm requires having a reference dataset (here, the three London scans) against which to compare the unknown dataset (the 25 participants in Auckland). It therefore depends on how likely it is that two comparable datasets are acquired at different sites or in independent projects. However, this scenario is very realistic: retinotopic mapping experiments are relatively standardized procedures used in many studies. Moreover, since we observed some of the best reidentification accuracy for coarse-scale activation maps, it is highly plausible that similar results could be found for simple localizer, activation, or resting-state functional connectivity studies.

In conclusion, sharing neuroimaging data is crucial for transparent and reproducible research practices, and generation of new knowledge that exceeds the original project aims. However, our present work highlights the need for appropriate safeguards against possible data misuse, such as data matching, that may allow reidentification of participants data.

## Data availability

The data and MATLAB analysis code necessary for reproducing these findings are available at osf.io/u9qn4. The data matrix has been shuffled to break the link between data and anatomical location within the template brain. It is therefore not possible to plot the maps from these data.

